# SeedMeasure: an efficient approach and open-source program to quantify seed size

**DOI:** 10.64898/2026.06.27.734974

**Authors:** Benjamin Sims, Allison Gaudinier, Benjamin K. Blackman

## Abstract

**Premise:** Seed size and morphology are critical traits in agriculture, ecology, and genetics, but high-throughput quantification of these traits is often limited by labor-intensive manual measurements or expensive, platform-specific imaging software.

**Methods and Results:** We developed SeedMeasure, a lightweight, open-source, and cross-platform command-line tool written in Python that automates the measurement of seed area, length, and width from images. Using a simple imaging setup, the program processes images by correcting for perspective skew, filtering debris, and exports quantitative data alongside quality-check images.

We validated SeedMeasure across nine diverse species, ranging from small *Arabidopsis thaliana* seeds to large *Zea mays* kernels. The tool quickly handles images using multithreading and demonstrates high reproducibility, yielding low coefficients of variation across repeated runs.

**Conclusions:** Compared to existing software, SeedMeasure is free, offers faster processing through parallel computing, and provides standalone executables that require no programming dependencies. SeedMeasure offers an accessible, cost-effective, and high-throughput approach for rapid phenotypic profiling, making advanced seed morphological analysis available to researchers without specialized laboratory hardware.

## INTRODUCTION

Seed size is a trait with applications across broad areas of plant biology. 60% of the world’s caloric needs are met by five seed crops: rice (*Oryza sativa*), wheat (*Triticum aestivum*), maize (*Zea mays*), sorghum (*Sorghum bicolor*), and millets. (FAO, 2018). For many crop species, seed size is a direct measure of yield and grain quality, making high-throughput screening essential to crop improvement programs. Beyond agricultural applications, quantifying seed size in a wide range of species with varying sizes, colors, and transparency is important for ecological research (e.g., Moles et al., 2007), seed banking (e.g., Probert et al., 2009), environmental response phenotyping (e.g., Donohue et al., 2009, Imbert & Ronce, 2001), genetic studies of natural trait variation (e.g., Tanabata et al., 2012), and developmental genetic screens (e.g., Herridge et al., 2011). Here, we describe SeedMeasure, a freely available and easy-to-use program that allows users to accurately and efficiently quantify seed size and morphology across a diversity of plant species.

Seed dimensions are commonly measured by length, width, and cross-sectional area. Manually taking seed length and width measurements with calipers for many grain crops is straightforward, but tedious and time consuming. Therefore, quantifying seed size for large volumes of seed is often done indirectly by measuring 1000-grain weight (Wang et al., 2023), but this strategy either requires specialized equipment or labor intensive procedures to count out 1000-seed aliquots. For species with smaller seeds, manual measurement of seed size is nearly impossible. Seeds from important model species, such as *Arabidopsis thaliana*, are often just a fraction of a millimeter long. Computer vision-based methods for measuring seeds offer a faster, more consistent approach for quantifying seed traits across large sample sets. However, many existing imaging programs are platform-specific, no longer supported, or complex to use (Zhu et al., 2021; Whan et al., 2014; Tanabata et al., 2012).

Here, we describe SeedMeasure, a lightweight, open-source command line tool for automated measurement of seed size from high resolution digital images. The program operates across operating systems, only requires a minimal imaging setup, and outputs quantitative data for downstream analysis. We present the imaging protocol, program workflow, and performance of SeedMeasure, and demonstrate its accuracy and reproducibility across multiple plant species of both agricultural and ecological importance.

## METHODS AND RESULTS

### Seed used in pipeline development

We used several seed lines maintained in the Blackman Lab at UC Berkeley, including *Arabidopsis thaliana* Col-0; *Helianthus annuus* HA243 (USDA GRIN ID: PI 650594), HA412 (USDA GRIN ID: PI 603993); *Mimulus guttatus* MAC inbred line seeds (Sacramento County, CA; 38° 14’ 56” N, 121° 05’ 26” W); and *Nicotiana benthamiana*. Additional seeds were obtained from other labs at UC Berkeley. Seeds of the *Sorghum bicolor* RTx430 and *Zea mays* B73 genotypes were supplied by the Lemaux Lab. The Krasileva Lab at UC Berkeley supplied *Triticum aestivum* seeds. The Savage Lab at UC Berkeley supplied *Oryza sativa* Kitaake seeds. We purchased *Penstemon clutei* seeds from PlantFlowerSeeds on Etsy.com.

### Seed imaging

We printed a 100 × 100 mm reference box on a sheet of white inkjet photo paper as the imaging background. This paper has a fine, uniform texture when backlit and is ideal for consistent measurements with SeedMeasure. We recommend measuring the external dimensions of your reference box after printing to control for variation in printing scale. For larger seeds, we imaged only those that were filled and looked viable, but this was not possible for small seeds. We spread out seeds evenly within the printed box to avoid clumping or overlapping, cleaned non-seed debris, and imaged the seeds over a light box (MPN E-21343) with a Samsung Galaxy S23 cellphone camera using at 50 MP, ISO 50, and a shutter speed of 1/50 s, with all other settings on auto. To minimize shadows and ensure maximum contrast between seeds and the background, we placed a 15 cm tall cardboard over the seeds with a 1 x 2 cm hole in the top for the cellphone camera lens (Figure 1).

**Figure 1.**
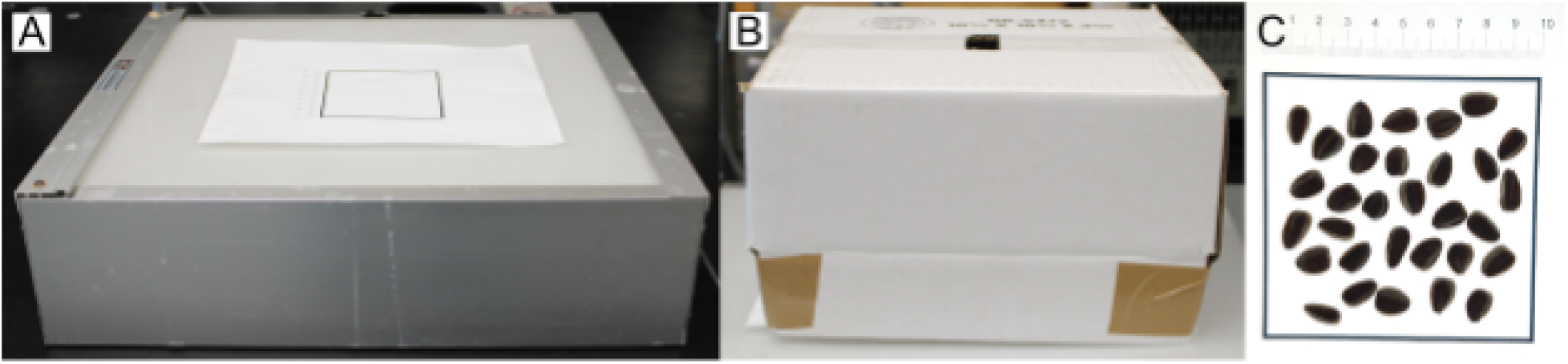
Seed imaging setup. We placed evenly spaced seeds on top of a printed 10 by 10 cm box (C) on top of an x-ray light box (A) to increase contrast between the seeds and background. We used a cardboard box (B) to exclude light and to shoot our photos at a consistent distance and angle.

### Image processing and measurement

After receiving the image as input, the program identifies the reference box in each image and uses it to correct for skew. Images taken at oblique angles are corrected back to a perfect square before the seeds are measured. This process allows for photos to be taken without requiring a tripod setup, as the cardboard box maintains lighting and camera working distance consistency. Using a consistent image setup allows for easier image processing and ensures that common program parameters can be used across an experiment. We took each image with a sample label beside the printed box to ensure our photos were not mislabeled. The box we printed (Figure 1C) contains a ruler along its exterior as a reference, but this is not seen by the program which crops out all regions outside the box without interference in analysis. The program excludes seeds located too close to the border of the box (within 10% of the edge) to minimize measurement error due to out-of-focus boundaries, but the user can change or remove this margin. Then, the program measures each remaining seed individually. For every seed detected, the program records cross-sectional area, width, length and the seed count per image. Width and length are calculated as the small and large dimensions respectively of a minimum-area bounding box drawn around a seed. Each measurement is written to a comma-separated values (CSV) file for downstream analysis (Figure 2).

**Figure 2.**
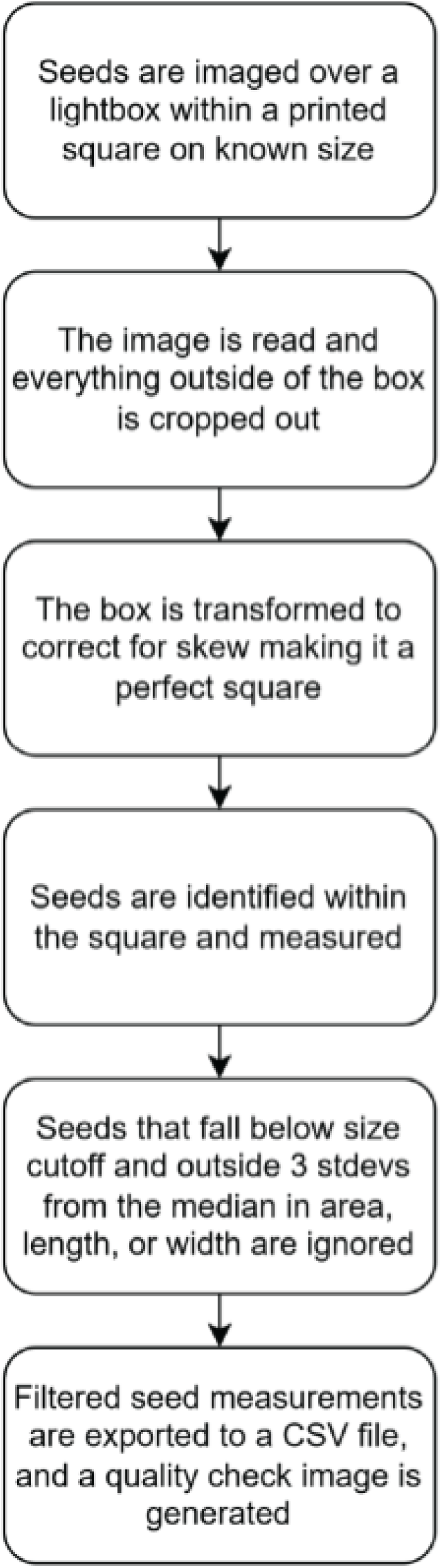
Overview of SeedMeasure’s image processing workflow. Upon receiving an image, SeedMeasure detects the bounding box, finds seeds, and outputs measurements to a CSV file.

To verify accuracy, the program also produces a quality check image with measured regions highlighted in red. This visualization provides a rapid visual assessment of whether the program identified seeds correctly and whether debris or artifacts are mistakenly measured.

### Program settings and automated filtering

To minimize inclusion of debris or clumped seeds, the program filters out objects whose area, length, width, or length-to-width ratio fall more than three standard deviations from the median of the sample. These objects are discarded from the dataset.

Seed characteristics vary across species, requiring different image analysis settings. Semi-translucent seeds, including *Z. mays* and *T. aestivum*, required a lower thresholding constant (inverse to the effective light sensitivity of the program) of 1 and 15, respectively, while all other seeds used a constant of 20. Seeds of different sizes also required minimum area cutoffs slightly smaller than the smallest seed in each set. We used cutoffs of 0.0, 0.0, and 0.1 mm² for the smallest seeds (*A. thaliana*, *M. guttatus*, and *N. benthamiana*, respectively), 0.1, 0.2, 1.0, and 1.0 mm² for intermediate seeds (*P. clutei*, *T. aestivum*, *O. sativa*, and *S. bicolor*), and 10 mm² for the much larger seeds of *H. annuus* and *Z. mays*. All other settings were kept consistent across seed types. Please refer to the supplemental how-to guide for detailed troubleshooting instructions.

### System requirements

SeedMeasure is a command line utility written in Python and built with NumPy (Harris et al., 2020) and OpenCV (Bardski, 2000). SeedMeasure will run on any device with at least Python 3.6, OpenCV 3.0, and NumPy 1.13 to run; however, we strongly recommend upgrading these packages to at least versions 3.8, 4.0, and 1.14 respectively. Alternatively, we provide standalone executables of the script compiled for Windows (x86-64), Mac (x86-64 and Apple Silicon), and Linux (x86-64), which can be run as-is on modern computer hardware and do not require the installation of any dependencies.

### Performance and reproducibility

The program processes each image in approximately 1 second on a laptop with an 8 core AMD Ryzen 7 7840HS processor running Windows 11. We supplied seeds that required the same settings as batches and enabled multithreading allowing SeedMeasure to process 8 images concurrently using 1 CPU per image. We verified accuracy and reproducibility by reshuffling and reimaging seed samples 8 times each for *O. sativa*, *T. aestivum*, and *M. guttatus* which produced consistent seed size distributions across all replicates (Figure 4).

### Seed size distributions across species

SeedMeasure’s versatility allows for the measurement of seeds of many sizes across multiple species (Figure 3; Table 1). Analysis of a wide variety of seeds make SeedMeasure suitable for a large number of applications, from measuring the size of grain to assess crop yields to counting seeds of native plant species used in seed banking.

**Figure 3.**
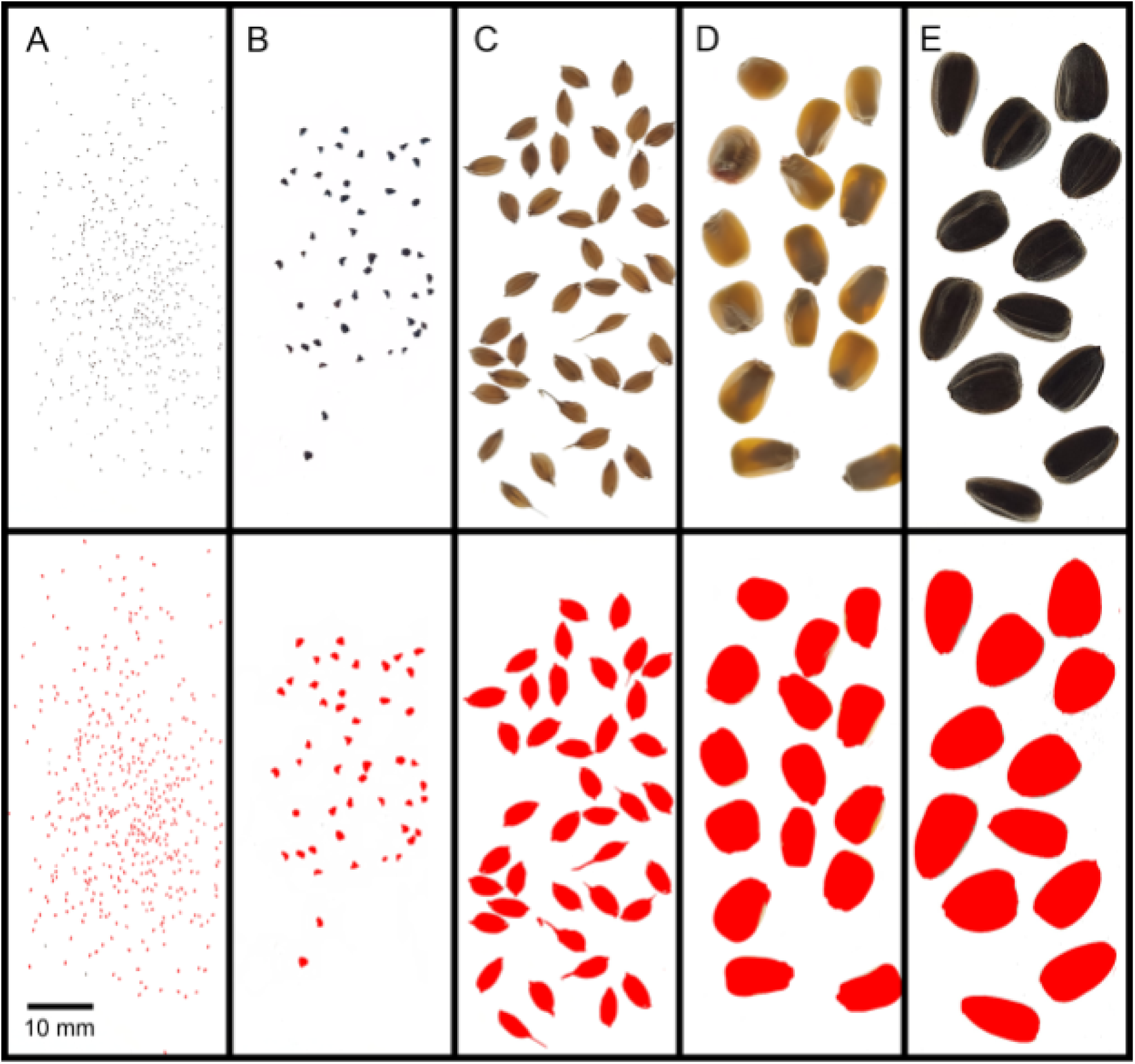
Snapshots of images of *M. guttatus* (A), *P. cluteiI* (B), *T. aestivum* (C), *Z. mays* (D), and *H. annuus* (E) supplied to SeedMeasure (top row), paired with the each image’s auto-generated quality check image file showing the seeds SeedMeasure visualized (bottom row).

**Table 1.** Mean seed sizes for each of the 10 lines measured including 1 standard deviation from the mean.

| Species | Mean Area<br>(mm<sup>2</sup>) | Length (mm) | Width (mm) | Number of<br>Seeds in<br>Sample |
| --- | --- | --- | --- | --- |
| *A. thaliana* Col-0 | 0.11 ± 0.02 | 0.46 ± 0.04 | 0.31 ± 0.03 | 292 |
| *H. annuus*<br>HA243 | 30.00 ± 10.15 | 102.51 ± 1.30 | 14.08 ± 0.98 | 30 |
| *H. annuus*<br>HA412 | 52.09 ± 9.15 | 11.62 ± 0.52 | 5.92 ± 0.91 | 58 |
| *M. guttatus* MAC | 0.12 ± 0.02 | 0.52 ± 0.05 | 0.30 ± 0.04 | 512 |
| *N. benthamiana* | 0.32 ± 0.05 | 0.78 ± 0.10 | 0.53 ± 0.06 | 66 |
| *O. sativa*<br>Kitaake | 16.64 ± 1.71 | 7.31 ± 0.97 | 3.35 ± 0.24 | 189 |
| *P. clutei* | 1.43 ± 0.36 | 1.66 ± 0.22 | 1.23 ± 0.19 | 84 |
| *S. bicolor*<br>RTx430 | 15.26 ± 1.12 | 4.44 ± 0.18 | 4.31 ± 0.16 | 100 |
| *T. aestivum* | 16.25 ± 2.57 | 6.55 ± 0.40 | 3.27 ± 0.37 | 113 |
| *Z. mays* B73 | 63.13 ± 7.95 | 11.37 ± 0.85 | 7.63 ± 0.83 | 39 |

### Consistency between runs

SeedMeasure produces consistent measurements given the same set of seeds. When we reshuffled and measured the same samples of *M. guttatus*, *O. sativa*, and *T. aestivum* seeds 8 times each, SeedMeasure produced very similar seed area distributions with coefficients of variation for the mean area of each sample at 2.41%, 1.86%, 0.95% respectively (Figure 4). Seeds will often fall in different orientations after each reshuffling resulting in slightly different measurements between runs. Seed clumping and resulting variation in the number of seeds automatically filtered from the count may also contribute to this variation.

**Figure 4.**
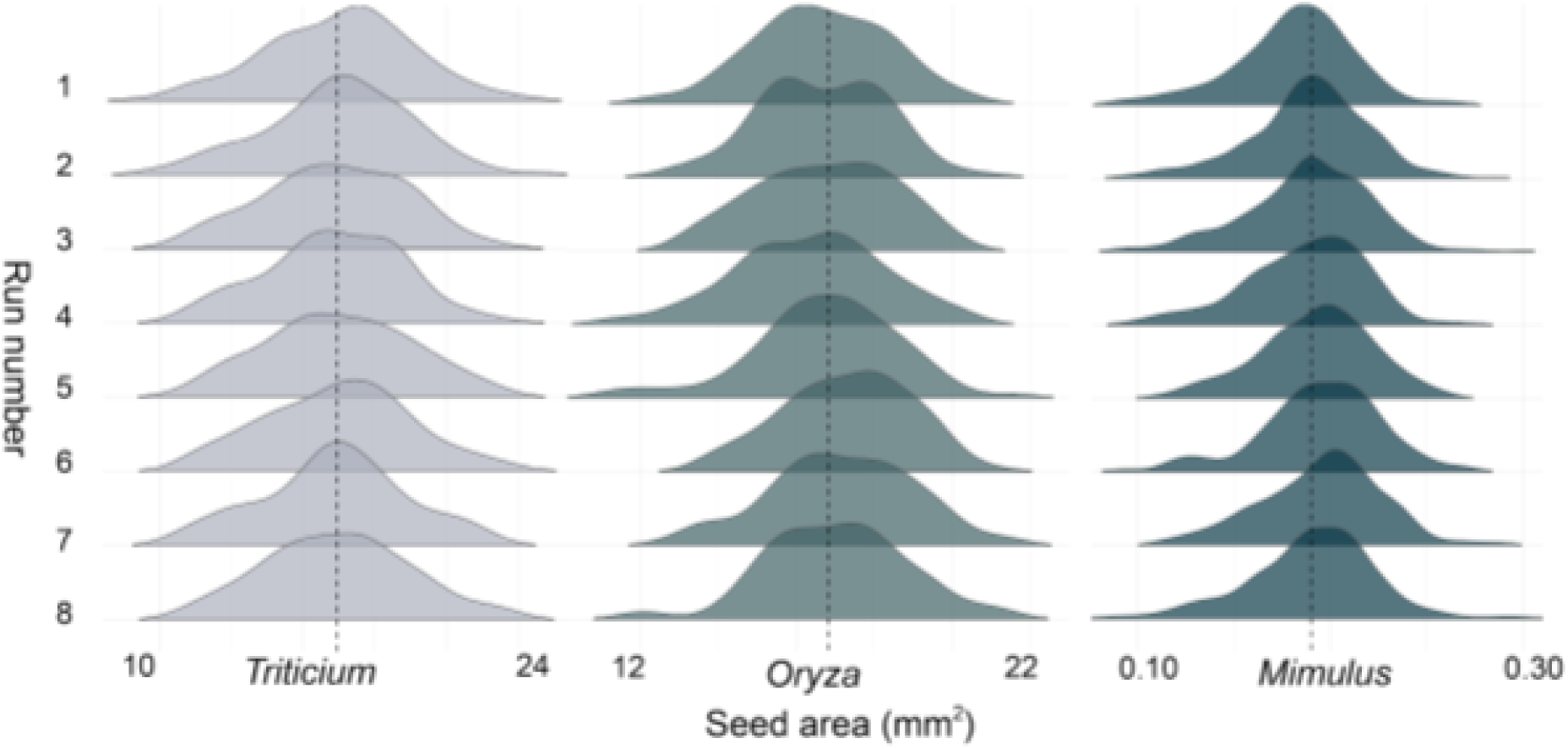
Seed measurement consistency across 8 runs of *T. aestivum* (left), *O. sativa* (center), and *M. guttatus* (right). Each density plot represents a separate measurement of seed area in the same sample of seeds.

### Performance and reproducibility

Compared to similar programs, SeedMeasure is free to use, quantifies seed area and shape the fastest, and runs on the widest variety of computer operating systems (Zhu et al., 2021; Whan et al., 2014; Tanabata et al., 2012). Unlike similar programs, SeedMeasure’s use of multiprocessing allows the script to take full advantage of the multiple logical processors in modern CPUs, scaling performance linearly with the number of logical processors available on your system. SeedMeasure’s Python script is compatible with any operating system with Python installed, and standalone executables are available for Mac, Windows, and Linux. Additionally, the modularity of SeedMeasure’s code enables anyone familiar with Python to create and add custom features to suit their specific needs.

## CONCLUSIONS

SeedMeasure provides a fast, high-throughput method to quantify the area, length, and width of seeds across diverse species. The program is compatible with most common operating systems and requires minimal setup. Furthermore, the images analyzed by SeedMeasure require basic equipment commonly found in a research laboratory to generate high precision measurements. They can be captured with a handheld camera or smartphone using a simple backlit sheet of paper. This versatility makes the tool highly adaptable for a wide range of applications, from ecological and crop studies focusing on morphology and grain yield to genetic research requiring precise phenotypic data. Altogether, these features make SeedMeasure an accessible, cost-effective solution for researchers seeking to streamline seed quantification without specialized laboratory hardware.

## Acknowledgements

This work was supported by funding from the National Science Foundation (IOS-2222464) and the United States Department of Agriculture (USDA 58-3060-4-011) to B. K. Blackman. The authors thank the Lemaux, Krasileva, and Savage labs for providing seed of various species for evaluation of our method and software.

## AUTHOR CONTRIBUTIONS

B. Sims contributed to the conceptualization, methodology, software development and distribution, data curation and analysis, and manuscript writing. A. Gaudinier contributed to the conceptualization, methodology, data analysis, supervision, and manuscript writing. B. K. Blackman contributed to the conceptualization, supervision, project administration, funding acquisition, and manuscript writing.

## DATA AVAILABILITY

SeedMeasure is open sourced (MIT License) and is available as both a Python file and as standalone executables, which require no dependencies. All source code and sample images are available for download on Github https://doi.org/10.5281/zenodo.20752879.

## APPENDIX

### Seed Imaging How-To Guide

Seeds must be imaged against a white sheet of photo paper within a printed reference box. While standard printer paper can be used, its grainier texture when backlit could cause SeedMeasure to mistake the background for seeds, especially when those seeds are translucent or pale-colored. The printed box can be any size depending on your needs, but it must be a perfect square. Its outside dimensions are supplied (which SeedMeasure assumes to be in millimeters) as an argument using the -w flag. The paper used for seed imaging should be placed on a lightbox in a darkened room to minimize shadows and ensure maximum contrast between seeds and background. Seeds should be spread apart manually before imaging to reduce clumping. SeedMeasure will exclude seeds that appear to be clumped, but minimizing the number of clumped seeds manually will improve accuracy. Images can be captured with a smartphone camera (ideally having a 12 MP sensor for medium to large-sized seeds, or a 50 MP sensor for Arabidopsis thaliana sized seeds or smaller).

The filename of each input image will be used as the line name for measurements in the output CSV file (e.g., aribidopsis_101.jpg is recorded as arabidopsis_101 in line_name column for each row of the output). You can supply these images individually to the program using the -i flag, or you can put all your images in the same directory to analyze them in a single batch.

### Image Processing and Measurement

SeedMeasure detects the seed bounding box by searching for the largest square (by area) in your image. Make sure any labels and seed packets you image are smaller than the bounding box.

Seeds located too close to the border of the box (within 5% of the edge by default) are excluded to minimize measurement error due to out-of-focus boundaries, but the user can change or remove this margin by supplying a percentage of the total box width as the -m flag. After every seed is detected, SeedMeasure records cross-sectional area, length (the maximum distance between two points on the seed’s perimeter), and width (the minimum distance between two points on the seed’s perimeter). Each measurement is written to a comma-separated values (CSV) file for downstream analysis with the total seed count per image recorded in its own column.

To verify accuracy, SeedMeasure also produces a quality check (QC) image with measured seeds highlighted in red while all other objects remain gray.

### Automated Filtering

SeedMeasure automatically filters out objects whose area, length, width, or length-to-width ratio fall more than three standard deviations from the median of the sample. These objects are discarded from the dataset. Automated filtering may not be appropriate if your seeds vary greatly in size or shape. Additionally, excess debris in images will result in less accurate filtering. Users may optionally disable or change the number of standard deviations used for this filtering feature with the -f flag.

### Performance and Reproducibility

When supplying a directory of many images at once, users can supply the number of threads to use with the -t flag, allowing SeedMeasure to process multiple images concurrently. The speed of image analysis scales proportionally to the number of threads available.

### Troubleshooting

Images taken at varying resolutions and brightnesses may require fine tuning of Seed Measurer’s thresholding settings. Using default settings, users may find that SeedMeasure’s thresholding is too sensitive, which will often result in details from the imaging background’s texture registering as seeds in the QC image. To fix this, try increasing the value given to the -k flag in increments of 1 until the background is no longer highlighted in red in your QC images. If these issues persist, try providing a minimum area cutoff using the -c flag set to a value just below the expected minimum cross-sectional area of your seed.

If SeedMeasure detects no seeds in your image, try decreasing the constant until your seeds show up in the QC image. Filtering settings may also help if extra large, small, or oddly shaped seeds are flagged as outliers and excluded from analysis. Supplying a lower filtering level using the -f flag or setting filtering to 0 to disable it completely may resolve these issues. If clumped seeds or debris are not excluded from analysis, try increasing the filtering level or manually separate your seeds, remove excess debris, and re-take your image.

## Notes

https://github.com/adequate-soup/seed-measure

### Competing Interest Statement

The authors have declared no competing interest.

